# Relative importance of bacterivorous mixotrophs in an estuary-coast environment

**DOI:** 10.1101/2023.09.30.560290

**Authors:** Qian Li, Kaiyi Dong, Ying Wang, Kyle F. Edwards

## Abstract

Mixotrophic eukaryotes are important bacterivores in oligotrophic open oceans, but their significance as grazers in more nutrient-rich waters is less clear. Here we investigated the bacterivory partition between mixotrophs and heterotrophs in a productive, estuary-influenced coastal region in East China Sea. We found ubiquitous, actively feeding phytoplankton populations and taxa with mixotrophic potential, by identifying ingestion of fluorescent prey surrogate and analyzing community 18S rRNA gene amplicons. Potential and active mixotrophs accounted for 10-63% of total eukaryotic community and 17-69% of bacterivores observed, respectively, contributing 6-48% of estimated in situ bacterivory. The much higher mixotroph fitness outside of the turbid plume were potentially driven by increased light and decreased nutrients availability. Our results suggest that, although heterotrophs dominated overall in situ bacterivory, mixotrophs were abundant and important bacterivores in this low-latitude mesotrophic coastal region.

**Scientific Significance Statement:** Mixotrophy, the combing of photosynthetic and phagotrophic nutrition, can dramatically increase primary production and alter material movements in aquatic food webs. Understanding the distribution and fitness of phagotrophic mixotrophs in a range of habitats are therefore critical to properly understand the biogeochemical cycles of the ocean. We developed a robust method to estimate active mixotrophs in situ and demonstrate rapid shifts of bacterivory partition between mixotrophs and heterotrophs across an estuary-coast environment in East China Sea. Our results help fill the knowledge gap of mixotroph niches in mesotrophic coastal environments that are impacted by riverine discharge, shedding light on relationships among mixotrophy, light and nutrient conditions.

## Introduction

Heterotrophic flagellates and microzooplankton were traditionally considered to be the primary, if not sole grazers of bacteria in aquatic food webs, but now there has been increasing recognition of the importance of unicellular phagotrophic mixotrophs (i.e. planktonic eukaryotes that are capable of photosynthesis and phagocytosis) (Sherr and Sherr, 2002; Stoecker et al., 2017). Model simulations suggest that mixotrophy can substantially increase primary production and energy transfer efficiency (Stoecker et al., 2009; Mitra et al., 2014; Ward and Follows, 2016), meanwhile bacterivory by mixotrophs may introduce different impacts on elemental flux and biogeochemical cycling. For instance, heterotrophs seem to respire more of the organic carbon and other elements from ingested prey to acquire energy (Hansen et al., 1997) whereas mixotrophs may utilize prey biomass primarily for nutrients, subsidizing/facilitating energy and organic carbon production through photosynthesis (Caron et al., 1993; Ptacnik et al., 2016; Mitra and Flynn, 2023).

The relative prevalence and success of mixotrophs has been commonly associated with oligotrophic environments (Unrein et al., 2007; Hartmann et al., 2012; Duhamel et al., 2019; Li et al., 2022), suggesting that low nutrients may select for mixotrophy (Lindehoff et al. 2010; Edwards et al., 2019). However, mixotrophs and mixotrophic bacterivory have also been reported to be ubiquitous in systems with relatively high nutrients supply, such as equatorial east Pacific (high light) (Stukel et al., 2011) and temperate estuaries adjacent to north Atlantic (low light) (Millette et al., 2017; 2021). Other studies have found higher irradiance to benefit mixotrophs, presumably because their phagotrophic abilities come at the expense of phototrophic performance (Raven 1997; Tittel et al., 2003; Edwards et al., 2022). These varied results coupled with the observed functional diversity (Li et al., 2022) and metabolic plasticity among mixotrophs (Li et al., 2021), illustrate that there are multiple interacting variables that determine whether mixotrophs will have an ecological advantage in a particular habitat. Understanding the range of habitats and mechanisms allow mixotrophs to thrive is therefore critical for interpreting their ecosystem-level biogeochemical consequences, yet remain challenging (Leles et al., 2021).

In this study we ask what are the major mixotrophic populations, how fast can they graze and what are the factors possibly affect bacterivory in an estuary-coastal region that receives freshwater discharge from the Yangtze River on the east coast of China. Hypotheses were proposed to see a shift between mixotrophic and heterotrophic grazer communities and bacterivory contributions along that estuary-coast environmental gradient, corresponding to the changing inorganic nutrients and other biophysical conditions. To answer these questions and testify our hypotheses, we developed a robust method to estimate active mixotrophs in situ via tracer experiments with fluorescent prey surrogates coupled with epifluorescence microscopic observation and high throughput flow cytometry screening. We also used 18S rRNA gene metabarcoding to evaluate overall eukaryotic taxonomy and identify potential mixotrophic bacterivores. Collectively, we aimed to investigate whether a representative coastal region influenced by river discharge favors mixotrophs.

## Materials and Methods

### Study sites and biochemical parameters

Water samples were collected during Nov 8-15, 2021 on board the ‘Zhe Yu Ke’ research vessel, from one estuary station and three shelf stations in the coastal East China Sea (ECS). Surface water from 2 m depth was collected by rosette mounted on a SBE19plus CTD equipped with auxiliary sensors. Inorganic nutrients including DIN (NO_3_) and DIP (PO_4_) were measured in the lab, using the Auto Discrete Chemical Analyzer (Smartchem 600, AMS Alliance, Italy). Total chlorophyll a was filtered onto 0.7 µm Whatman GF/F glass fiber filters, extracted with acetone overnight in the dark and measured on a Turner Designs AU-10 fluorometer. Cell abundance samples for eukaryotes and bacteria were fixed with glutaraldehyde at 0.5% final concentrations (Sigma Aldrich, USA) and stored at -80 ℃ until analyzed via flow cytometry in the lab (CytoFLEX S, Beckman Coulter, USA).

### Short-term grazing experiments

Three prey surrogates were first assessed prior to the cruise to identify the most suitable tracers, among which the 1.0 µm green-yellow fluorescent beads produced the best detection results by flow cytometry (Supplementary Fig. S1), and were thus chosen for the grazing experiments. During the cruise, approximately 200 mL pre-filtered (200 µm nylon mesh) seawater in triplicate was amended with 1.0 µm beads at a final concentration of ∼2.5×10^5^ mL^-1^, accounting for 20-50% of the in situ bacteria concentrations. Samples were incubated for one hour under 14-18℃, ∼70 µE sec^-1^ m^-2^, and 12h:12h light/dark cycle. Twenty-mL fixed (0.5% glutaraldehyde) and stained aliquots (0.1 µg mL^-1^ DAPI; Biotium; USA) were filtered onto 2 µm polycarbonate membrane filters (Millipore) for microscopic examination (Olympus BX61 inverted microscopy).

### Active bacterivores identification

Under microscope, cells with ingested beads were counted as active bacterivores, and mixotrophs were distinguished based on the presence of conspicuous chloroplasts (organelles with red autofluorescence). Potential non-constitutive mixotrophs with keptoplastic plastids were treated as heterotrophs as we could not verify their mixotrophic activities. Each sample was scanned for 130-150 fields to decrease counting errors (Lund et al., 1958). Active bacterivores were also identified on flow cytometry (CytoFLEX S, Beckman Coulter). Total pigmented eukaryotes and heterotrophic flagellates were first identified via red fluorescence (chlorophyll a), blue fluorescence (stained nucleic acid) and size scatters, after which active bacterivores were distinguished as the same eukaryotic population, but which had also acquired the characteristic yellow-green fluorescence of ingested beads (γ_em_∼ 505 nm).

### Grazing rates estimate

The average per capita ingestion rate of beads (IR*_i_*, beads cell^-1^ h^-1^) in a sample was calculated using equation:

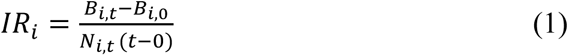

Where B_i,t_ and B_i,0_, are the total number of ingested beads observed at time t (i.e. one hour post incubation) and initial time point, of group *i* (bacterivores were binned into six size groups). N_i,t_ are the total active bacterivores observed at time t of group *i*.

The estimated in situ bacterivory rate (BR), i.e. the grazing impact of bacterivores in aggregate (bac. mL^-1^ d^-1^) were calculated using equations (2):

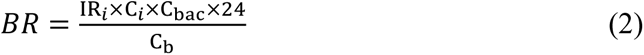

Where C*_i_*, C_bac_ and C_b_ are the concentrations (cells mL^-1^) of active bacterivores (group *i*), total bacteria, and fluorescent beads (initial concentrations), respectively. C*_i_* was calculated by multiplying mean cell counts per field under microscopy by 3.9×10^4^ fields (obtained by dividing the membrane filters’ area by each observational field’s area) and dividing by aliquot volume (mL).

### 18S rRNA gene metabarcoding

We vacuum filtered approximately 1 L seawater from each sample through 47 mm, 0.45 µm polycarbonate membrane filter (Millipore, USA) and extracted total environmental DNA which were used as template to amplify the hypervariable V4 region of 18S rRNA with primers 547F and V4R (Stoeck et al., 2010). PCR products from each sample were purified and pooled in equal amounts, followed by pair-end 2×250 bp sequencing using the Illumina NovaSeq platform at Shanghai Personal Biotechnology Co., Ltd, China. Quality check and chimeras removal of amplicon sequence variants (ASVs) were conducted using DADA2 (Callahan et al., 2016), which were further classified with two databases, the Protist Ribosomal Reference Database (Guillou et al., 2013) and SILVA (Pruesse et al., 2007) in QIIME2 (Bolyen et al., 2019). Classification disagreement between the two databases were manually curated by searching additional cultivated references using BLASTn and the NCBI GenBank (Supplementary Table S1). ASVs that were metazoa, freshwater/land-originated (mostly fungi and some amoeba at B3), or with low numbers were excluded from further analysis.

### Correction factors for 18S rRNA gene abundances

Ribosomal gene copy numbers vary substantially among eukaryotic groups, from as low as one copy in small flagellates to thousands of copies per cell in dinoflagellates (Zhu et al., 2005). This, along with biases in the amplification reaction (Bradley et al., 2016; Gonzalez et al. 2012), can result in relative gene abundances in amplicon pools that are significantly different from cell abundances (Karlusich et al., 2022). To correct for this bias, we followed the principle of a study by Martin et al. (2022) and developed our own empirical gene-to-cell correction factors (C.F.) for seven major groups that were distinguishable by morphology under microscopy (dinoflagellates, ciliates, radiolaria, cryptophytes, diatoms, other pigmented and heterotrophic eukaryotes), which were formulated as:

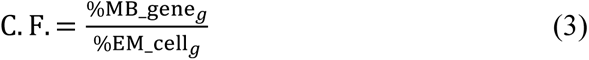

Where %MB_gene*_g_* and %EM_cell*_g_* are the relative abundances of group *g* derived from metabarcoding 18S rRNA reads, and from epifluorescence microscopy-based cell abundances, respectively. These C.F. were then averaged with those of Martin et al. (2022)’s to retrieve the final factors used in this study (results shown in Supplementary Fig. S2). The corrected relative abundance of group *g* (%MB_cell*_g_*) can be subsequently calculated using equation (4), and were used in all of our analysis except for Fig. 1b.

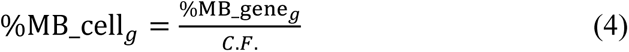

**Fig. 1.**
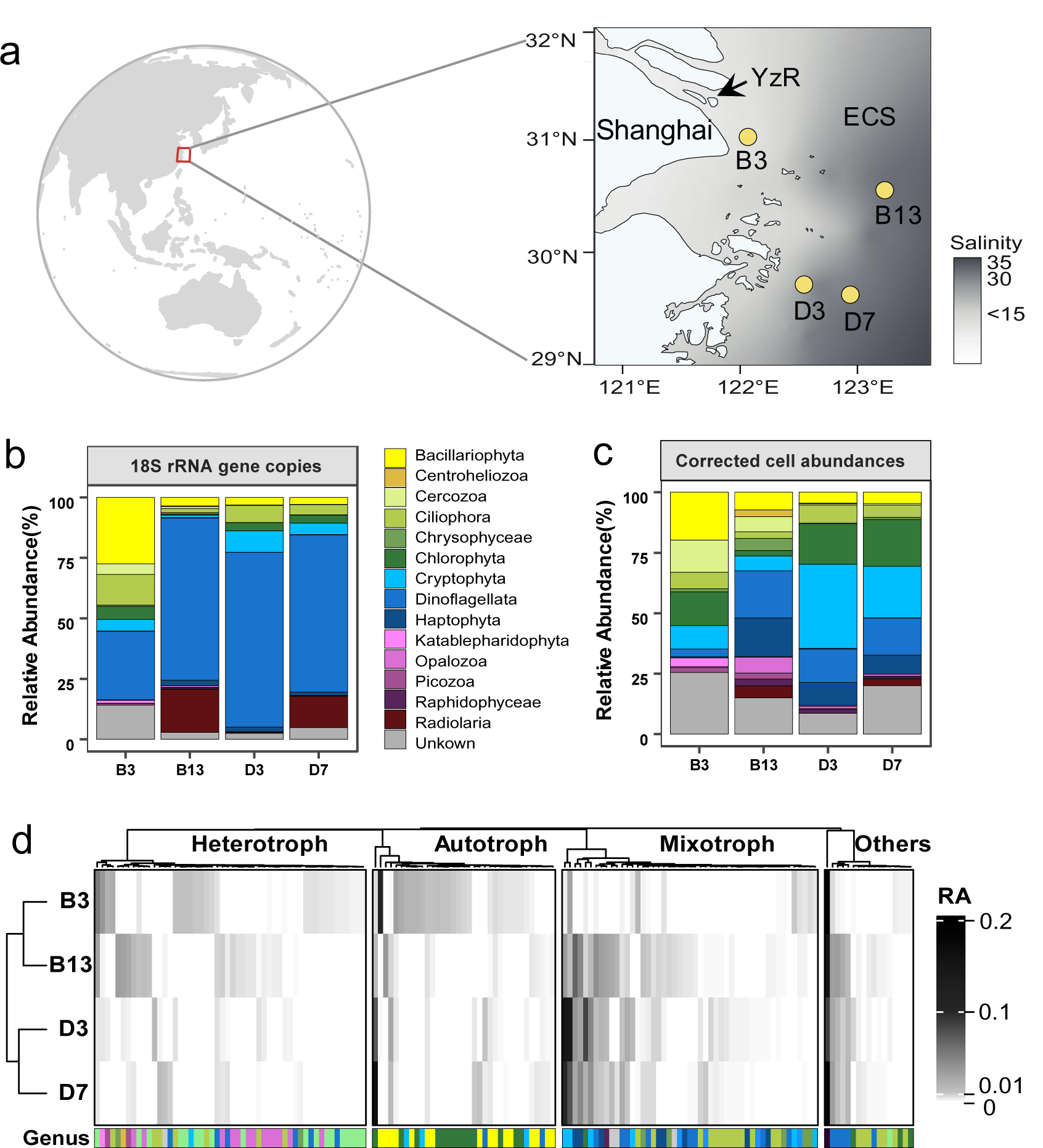
Oceanographic map with salinity in the background showing sampling sites of this study (**a**), 18S rRNA genes derived taxonomic composition of top twelve divisions before (**b**), and after gene-to-cells conversion using correction factors (**c**), and a heatmap constructed with Pearson distances showing trophic mode compositions in relative abundances (RA) using results in panel c but on genus level (**d**). Taxonomy in panels **b** and **c** were division levels based on the PR^2^ database ranks, with the exception that Ochrophyta was presented in three classes (Bacillariophyta, Chrysophyceae and Raphidophyceae). Note that unkown reads were not corrected for gene copies cell^-1^ which may result in an overestimated or underestimated percentage.

### Annotation of potential mixotrophs

We assigned each genus (or species when annotation was unavailable) to one of four nutritional categories, i.e. mixotrophy (including constitutive and non-constitutive mixotrophs), heterotrophy, autotrophy, or others, using evidence from synthesized databases (e.g. Faure et al., 2019; Mitra et al., 2023) and research articles (Supplementary Table S2). Note that one of the abundant genus *Micromonas* of Chlorophyta were assigned as autotrophs as evidence for phagotrophy was ambiguous (McKie-Krisberg and Sanders, 2014; Jimenez et al., 2021). Some Radiolaria and *Noctiluca* were considered as others/symbiotic, all Syndiniales were assigned as others/parasitic and unkown taxa or taxa with unavailable trophic evidence were all assigned as ‘others/unkown’.

### Statistical analysis

All of our statistical analysis and visualization were carried out in R, v3.6.3 (R Core Team, 2020). Pairwise t tests was performed using basic R functions. Redundancy analysis (RDA) and Spearman correlation matrix were applied to investigate significant correlations between mixotrophic fitness and biophysiochemical variables, using the vegan and psych packages for calculation and pheatmap package for visualization. ASV reads were first Hellinger-transformed and all independent variables were log transformed for normalization. Low Variance Inflation Factor (<10) were checked to eliminate co-linearity in the RDA.

## Results

### Hydrographic and biochemical characteristics of the study region

Geographically, the estuary station B3 within the plume region showed the lowest salinity (14.2) and temperature (14.2 °C) and the highest nutrients (62.1 µM NO_3_, 1.3 µM PO_4_) and turbidity (24.6 NTU), while the other stations (B13, D3, D7) exhibited high salinity (30-33) and temperature (18.7-19.5 °C), low turbidity (1.2-1.6 NTU) and relatively lower nutrients (8-17 µM NO_3_, 0.4-0.6 µM PO_4_) (Fig. 1a, Table 1). Therefore, the studied region is a typical estuary-to-shelf mesotrophic system that receives heavy riverine discharge resulting in a maximal turbid zone adjacent to station B3. Correspondingly, total Chl *a* were medium-high, ranging between 0.9-2.5 µg mL^-1^. Abundances of pigmented eukaryotes (1.2-1.9 × 10^4^ cells mL^-1^) were overall higher than heterotrophic flagellates (0.2-0.6 × 10^4^ cells mL^-1^), especially at stations D3 and D7. Heterotrophic bacteria and autotrophic bacteria (mostly *Synechococcus*) presented concentrations between 4.7-8.4 × 10^5^ cells mL^-1^ and 0.3-4.3 × 10^4^ cells mL^-1^, respectively (Table 1).

**Table 1.** Physical and biochemical characteristics of the study sites. Sta, Temp, Sal, Turb, PO_4_, NO_3_, TChl *a*, HBac, Syn, PE, HF stand for station, temperature, salinity, phosphate, nitrate, total chlorophyll a, heterotrophic bacteria, *Synechococcus*, pigmented flagellates, heterotrophic flagellates and potential mixotrophic flagellates, respectively.

| Sta | Temp<br>°C | Sal | Turb<br>NTU | PO<sub>4</sub><br>μM | NO<sub>3</sub><br>μM | TChl *a*<br>μg mL<sup>-1</sup> | HBac<br>cells mL<sup>-1</sup> | Syn<br>cells mL<sup>-1</sup> | PE<sup>a</sup><br>cells mL<sup>-1</sup> | HF<sup>a</sup><br>cells mL<sup>-1</sup> |
| --- | --- | --- | --- | --- | --- | --- | --- | --- | --- | --- |
| B3 | 14.2 | 14.2 | 24.6 | 1.3 | 62.1 | 1.2 | 4.7×10<sup>5</sup> | 3×10<sup>3</sup> | 1.5 (±0.5) ×10<sup>4</sup> | 6.1 (±1.7) ×10<sup>3</sup> |
| B13 | 19.4 | 33.5 | 1.2 | 0.6 | 9.5 | 0.9 | 6.9×10<sup>5</sup> | 2.8×10<sup>4</sup> | 1.2 (±0.4) ×10<sup>4</sup> | 3.6 (±1.4) ×10<sup>3</sup> |
| D3 | 18.7 | 30.5 | 2.0 | 0.6 | 16.2 | 2.3 | 10.4×10<sup>5</sup> | 4.3×10<sup>4</sup> | 1.7 (±0.3) ×10<sup>4</sup> | 1.9 (±0.7) ×10<sup>3</sup> |
| D7 | 19.5 | 32.7 | 1.1 | 0.4 | 8 | 2.5 | 8.4×10<sup>5</sup> | 3.5×10<sup>4</sup> | 1.9 (±0.5) ×10<sup>4</sup> | 1.9 (±0.5) ×10<sup>3</sup> |

<sup>a</sup>PE and HF abundance were the mean values averaged from flow cytometry and epifluorescent microscopy observations. Values in brackets are standard deviations between two methods.

### Metabarcoding-derived community composition and potential mixotrophs

Before the gene-to-cell correction (in %MB_gene), amplicon data suggested that all stations were dominated by 18S rRNA genes from Dinoflagellata, except for B3 which was co-dominated by Bacillariophyta and Dinoflagellata. Other abundant groups included Radiolaria (at B13 and D7) and Cryptophyta (at D3 and D7) (Fig. 1b). After correction for gene copy numbers (in %MB_cell) it is estimated that the three most abundant populations at B3 were Bacillariophyta, Chlorophyta and Cercozoa (Fig. 1c), among which the majority genus were annotated as autotrophs and heterotrophs (Fig. 1d; Supplementary Table S2). Station B13 was dominated by pigmented eukaryotes from Dinoflagellata and Haptophyta and heterotrophs from MAST group. Cryptophyta and Chlorophyta were both highly abundant at D3 and D7, followed by Dinoflagellata and Haptophyta (Fig. 1c). Among all pigmented eukaryotes from stations B13, D3 and D7, the majority of most abundant genera were potential mixotrophs, such as *Cryptomonadales* and *Teleaulax* of Cryptophyta, *Pyramimonas* of Chlorophyta, *Chrysochromulina* and *Phaeocystis* of Haptophyta, as well as *Gymnodinium* and *Heterocapsa* of Dinoflagellata (Fig. 1d; Supplementary Table S2). When considering both taxonomy and nutritional mode composition (in %MB_cell), the Pearson distance clustering suggested a closer association between B3 and B13, and D3 and D7, respectively (Fig. 1d). Particularly, the B transect had substantially more heterotrophs than the D transect (24-25% vs 5-7%), while B3 presented the lowest (potential) mixoptroph proportions among all four station.

### Microscopy and flow cytometry-identified active mixotrophs

Ubiquitous, actively-feeding mixotrophs and heterotrophs were identified from small (<5 µm), medium (5-10 µm), and large-sized (>10 µm) eukaryotes under epifluorescence microscopy (EM) (Fig. 2a). Among all six size categories, 5-10 µm heterotrophs and mixotrophs were the most abundant bacterivores, contributing a mean of 71.6% and 28.4% of total bacterivory across all stations (Supplementary Fig. S3). Averaged ingestion rates of all mixotrophs and heterotrophs varied from 1.5 to 3.8 beads cell^-1^ h^-1^, accompanied with estimated in situ abundances and bacterivory rates of 0.3-1.8 × 10^3^ cells mL^-1^ and 0.2-3.2 × 10^5^ bact. mL^-1^ d^-1^, respectively (Table 2).

**Fig. 2.**
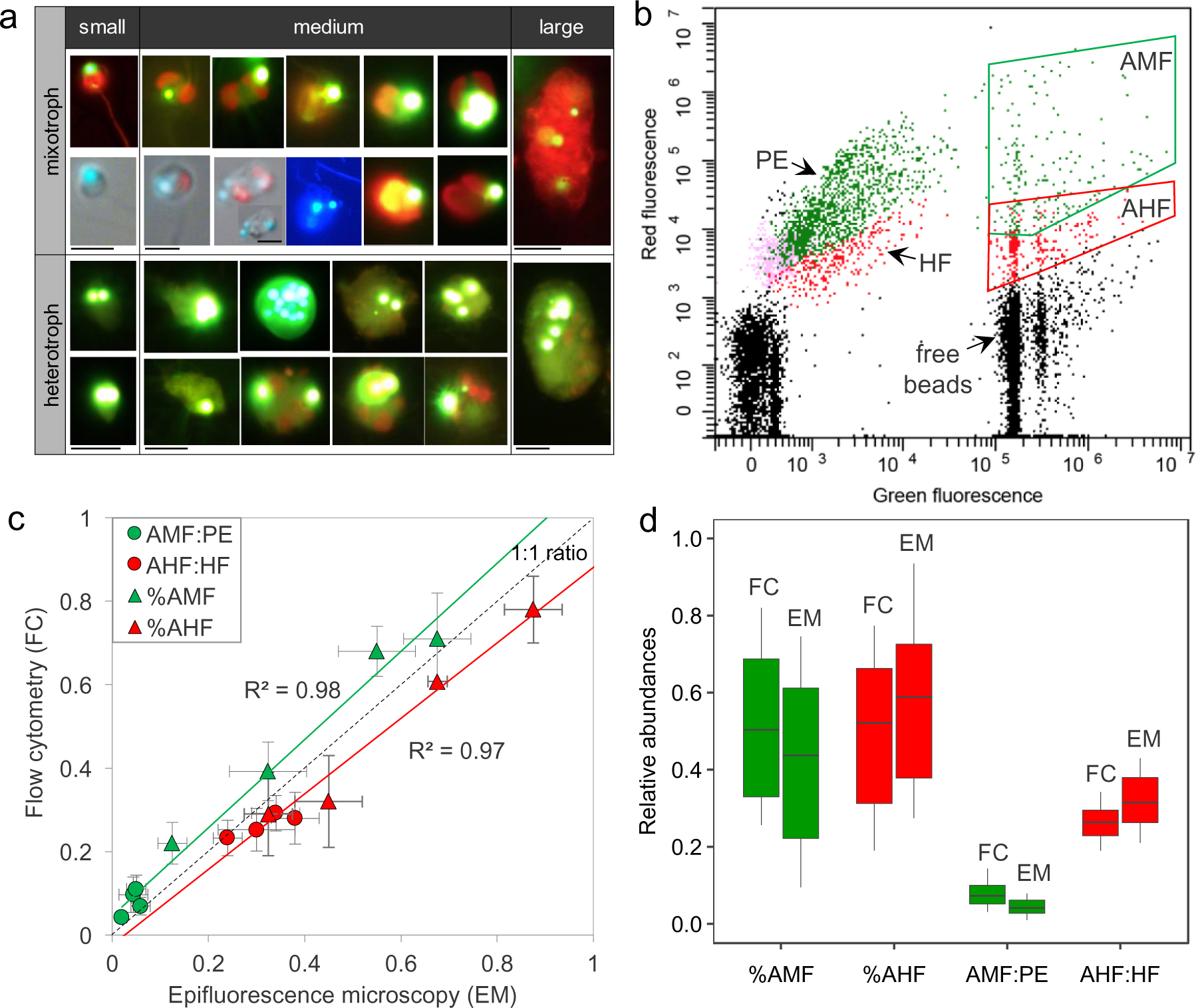
Epifluorescence microscopy images showing evidence of ingestion (beads in green-yellow, or cyan-blue fluorescence) by representative mixotrophs (chloroplasts in red or orange fluorescence) and heterotrophs among different size groups (**a**). Flow cytometric plot (**b**) from station B13 showing pigmented eukaryotes (PE) and active mixotrophs (AMF) in green dots, and heterotrophic flagellates (HF) and active heterotrophs (AHF) in red dots, initially distinguished by red fluorescence (chlorophyll *a*) and blue fluorescence (DAPI stained nucleic acid) as shown in Supplementary **Fig. S4**. Panel (**c**) compares results between epifluorescence microscopy (EM) and flow cytometry (FC) at each station, including relative abundances of AMF against PE, AHF against HF, as well as %AMF and %AHF against total bacterivores. Panel (**d**) demonstrate the same results in panel **c**, but with averaged values across all stations. All scale bars in panel **a** is 5 µm in length. The first three images in the lower mixotroph panel demonstrate the beads position in liquid samples (similar morphology), and the fourth and fifth images show the same cell (as in the upper panel) excited with blue (DAPI) and orange (phycobilins) fluorescence.

**Table 2.** Summary of grazing rates and abundances of mixotrophic and heterotrophic bacterivores in the investigated region. IR and BR denote ingestion rate (prey cell^-1^ h^-1^), and estimated in situ bacterivory rates (bact. consumed mL^-1^ d^-1^). Abbreviations of AMF/AHF and PMF denote actively feeding mixotrophic/heterotrophic flagellates and potential mixotrophic flagellates, respectively.

| Sta | IR_AMF<br>prey cell<sup>-1</sup> h<sup>-1</sup> | IR_AHF<br>prey cell<sup>-1</sup> h<sup>-1</sup> | PMF<sup>a</sup><br>cells mL<sup>-1</sup> | AMF<br>cells mL<sup>-1</sup> | AHF<br>cells mL<sup>-1</sup> | BR_AMF<br>bact. mL<sup>-1</sup> d<sup>-1</sup> | BR_AHF<br>bact. mL<sup>-1</sup> d<sup>-1</sup> |
| --- | --- | --- | --- | --- | --- | --- | --- |
| B3 | 1.7 | 3.8 | 1.4×10<sup>3</sup> | 2.6×10<sup>2</sup> | 1.8×10<sup>3</sup> | 2.0×10<sup>4</sup> | 3.2×10<sup>5</sup> |
| B13 | 1.8 | 3.5 | 4.6×10<sup>3</sup> | 5.9×10<sup>2</sup> | 1.2×10<sup>3</sup> | 6.9×10<sup>4</sup> | 2.8×10<sup>5</sup> |
| D3 | 1.5 | 3.4 | 1.1×10<sup>4</sup> | 9.4×10<sup>2</sup> | 4.5×10<sup>2</sup> | 1.4×10<sup>5</sup> | 1.5×10<sup>5</sup> |
| D7 | 1.8 | 3.3 | 8.1×10<sup>3</sup> | 8.8×10<sup>2</sup> | 7.2×10<sup>2</sup> | 1.3×10<sup>5</sup> | 1.9×10<sup>5</sup> |

<sup>a</sup>PMF abundances were estimated using mean pigmented eukaryotes abundances multiplying by 18S rRNA genes converted relative abundance of mixotrophs (out of total pigmented eukaryotes).

Flow cytometry (FC) approach was robust to identify active bacterivores (Fig. 2b; Supplementary Fig. S4). Nevertheless, relative abundances of active mixotrophs (against total pigmented eukaryotes) and heterotrophs (against total heterotrophic flagellates) revealed by FC were overall higher and lower compared to EM approach, i.e. 4-11% vs 2-6%, and 23-29% vs 24-38%, respectively (Fig. 2c-d). Proportions of mixotrophs (against total bacterivores) were 22-71% by FC method (mean value 50%), also higher than the EM-derived estimates of 13-68% (mean value 42%). Across stations, there were strong regression between results derived from two approaches, i.e. R^2^=0.98 for mixotrophs (in green line) and R^2^=0.97 for heterotrophs (in red line) (Fig. 2c).

### Mixotrophic fitness across the estuary-coast gradient

We evaluated the relative fitness of mixotrophs based on relative abundance of potential mixotrophs and proportion of active mixotrophs (averaged from both FC and EM estimates) at each station (Fig. 3a, b). Both fitness proxies increased from B3 (10% and 17%) to B13 (39% and 36%) and D7/D3 (41-63% and 62-69%), accompanied with dramatic decreases of turbidity and nutrients (e.g. DIN, DIP) (Table 1; Fig. 3a, b). Among selected variables (those passed variance inflation factor test of RDA), abundances of total pigmented eukaryotes were affiliated with potential mixotrophs (e.g. *Cryptomonadales*, *Teleaulax* and *Pyramimonas*) at stations D3/D7 on the RDA plot, directed away from turbidity and station B3. Strict autotrophs (e.g. *Skeletonema* and *Chaetoceros*) and heterotrophs (e.g. *Katablepharis* and Cercozoa_XXXX) were associated with B3 and high turbidity. B13 was separated from both B3 and D3/D7, associated with a mixture of mixotrophs (e.g. *Chrysochromulina* and *Heterocapsa*) and heterotrophs (e.g. MAST_3E and *Gyrodinium*) taxa (Fig. 3c, d).

**Fig. 3.**
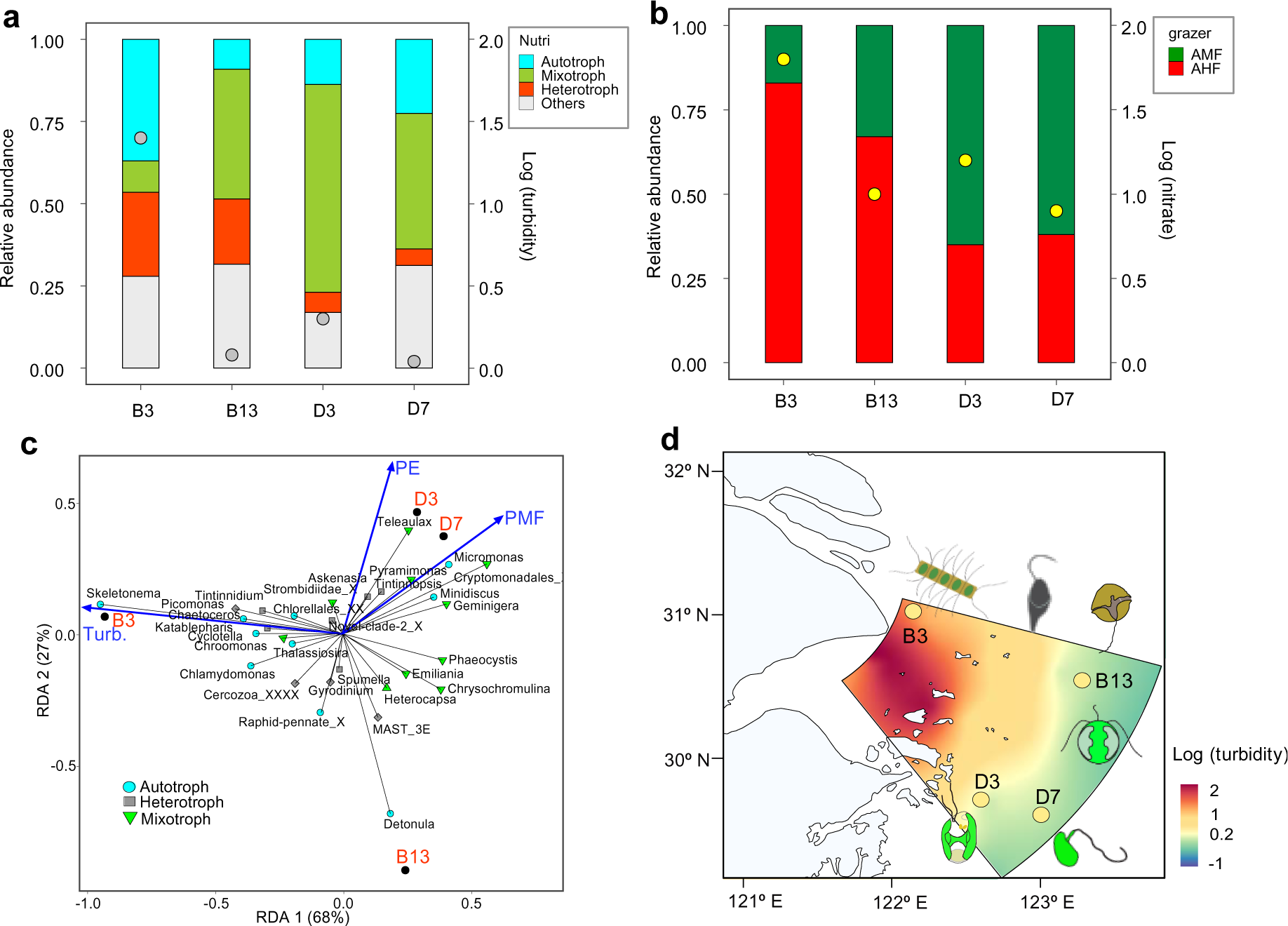
Composition of four nutritional modes annotated by 18S rRNA amplicon (**a**) and bacterivory partition between active mixotrophic flagellates (AMF) and heterotrophic flagellates (AHF) across stations (**b**). Log-transformed turbidity (NTU) in grey dots, and nitrate concentrations (µM) in yellow dots (both correspond to the second axes on the right) were overlay on top of bars in panel (a) and (b), respectively. Panel (**c**) shows the redundancy analysis (RDA) between the top 10 genera (summed across all stations) from each nutritional mode, and selected independent variables (i.e. PMF of potential mixotrophs, PE of pigmented eukaryotes and Turb of turbidity). Panel (**d**) is a turbidity map with representative bacterivore illustrations to demonstrate community shift among mixotrophs (green and brown colored flagellates), autotrophs (chained diatoms) and heterotrophs (grey colored flagellates). Taxa names with the “X” suffix mean that they could not be annotated on lower taxonomy ranks.

For the estimated bacterivory rates, mixotrophs from all size classes collectively contributed significantly higher proportions outside of the plume (20% at B13 and 41-48% at D3/D7) than B3 (6%) (Pair-wise student’s t test; P<0.05) (Supplementary Fig. S5a). The size category of dominant grazers did not vary across stations, i.e. all from the 5-10 µm medium size range, suggesting the prey (size) is likely controlling the grazer size and type rather than other environmental variables (Supplementary Fig. S5b, c). Comparison of other physical and biochemical conditions across stations revealed the highest similarity between D3 and D7, followed with B13 and then B3 (Supplementary Fig. S6a). Pearson correlation matrix suggested that bacterivory by mixotrophs were positively correlated with heterotrophic bacteria, *Synechoccoccus* and abundances of potential mixotrophs (P<0.05) (Supplementary Fig. S6b).

## Discussion

### Methodology to identify active mixotrophs

EM and FC approaches were demonstrated to be robust for retrieving active bacterivores from natural environments, but both have limits. Microscopic observation could provide direct phagocytosis evidence but were time-consuming and of low resolution. FC screening was high throughput but failed to estimate grazing rates. One challenge from using the EM method is to recognize the position of prey (inside or outside of the cell). Because many mixotrophic flagellates lack cell walls, i.e. shown as cells without outlines/borders under fluorescence excitation, sometimes prey appear to be, but may not be, attaching to (outside of) the cells (Fig. 2a). We have noticed the same issue during one of our previous study using mixotrophic cultures (*Florenciella* sp. of Dictyochophyceae), and we partially solved the problem by staining cell membranes (CellBright 488, Biotum, USA) to confirm that oftentimes prey were indeed ingested or in the process of entering the membrane (Li et al., 2021). In this study, we examined additional liquid samples under bright field (have clearer cell outline) to confirm phagocytosis in common mixotrophs seen on membrane filters (Fig. 2a). Nevertheless, ingestions rates derived from EM should be annotated with caution. Other studies have also indicated that 1.0 µm beads tend to stuck in the cell membrane during the process of phagocytosis by their mixotrophic grazers (Millette et al., 2021).

FC method generated higher estimates of mixotrophs than microscopy for our samples, agreeing to a study using similar method of FC and acidotropic probes by Costa et al. (2022). It was either due to underestimated EM results and/or overestimated FC results, and if the latter, these overestimates were likely from omnivorous heterotrophs that ingested both pigmented eukaryotes and bacterial prey surrogates (some of the lowest panel images in Fig. 2a). Combined methods using various tools with a focus on single cells (such as single cell sorting and sequencing) were suggested to retrieve the most reliable data for future mixotrophy research (especially for taxonomic identification) (Beisner et al., 2019).

### Trophic mode assignment

Annotation of putative trophic modes on the basis of taxonomy is another effective approach to study mixotrophy but can sometimes lead to biased interpretations. For example, some studies have found interspecific and intraspecific variations of trophic mode from ciliates *Mesodinium* sp. (Johnson et al. 2016) and dinoflagellate *Karlodinium* sp. (Calbet et al., 2011). Although others have suggested mixotrophy tend to broadly exist among different species within the same genus, such as *Ochromonas* of Chrysophyceae (Wilken et al., 2019), *Florenciella* of Dictyochophyceae (Li et al., 2022) and *Phaeocystis* of Haptophyta (Koppelle et al., 2022). Future experiments need to be carried out to verify whether these trophic variations are intrinsic (evolutionary) or due to plasticity under different physiological and environmental conditions.

### Enhanced mixotrophic fitness outside of the turbid estuary

Mixotrophs compete inorganic nutrients with autotrophs and organic nutrients (prey) with their heterotrophic competitors. The relative fitness and niches partition among them under different habitats remain an important question to answer. We chose an environment covers light limiting eutrophic estuary and coastal sites with higher irradiance and less nutrients. Our results suggested that mixotrophs were suppressed at the highly turbid estuary where strict autotrophs and heterotrophs were superior competitors, but became dominant outside of the estuary plume. These niche shifts are consistent with many studies that have suggested mixotrophs benefit from low-nutrients and high-irradiance environments (Tittel et al., 2003; Stukel et al., 2011; Anderson et al., 2019; Edwards et al., 2019; 2022). One explanation for this is that mixotrophs are penalized for possessing dual apparatus of phototrophy and phagotrophy within the same cells, i.e. have poorer photosynthetic performance comparing to autotrophs and lower ingestion rates comparing to heterotrophs (Raven, 1997). With enhanced irradiance, decreased nutrients, and abundant bacterial prey (e.g. sites outside of the estuary), mixotrophs benefited from both inorganic and organic nutrients which can offset nutrients limitation and stimulate more growth of mixotrophs. Although mixotrophs may have lower ingestion rates than their heterotrophic competitors, phototrophy-derived carbon and energy (when light is unlimited) can increase the competitive ability of mixotrophs against heterotrophs (tend to be energy-limited), meanwhile suppressing prey to an equal or lower concentration and making them comparable or superior competitors (Rothhaupt, 1996; Fischer et al., 2016; Edwards et al., 2023).

Overall, this study demonstrated an ecological niche for phagotrophic mixotrophs in a low-latitude mesotrophic coastal region impacted by land riverine inputs, expanding our knowledge of mixotrophic distribution and niche fitness. Finally, the rapid shifts in partitioning of bacterivory between mixotrophs and heterotrophs happened at small geographic scales (e.g. less than 100 kilometers), reiterating the importance of high-frequency and high-resolution sampling efforts for future research.

## Acknowledgement

We thank the Observation and Research Station of Marine Ecosystem in the Yangtze River Estuary, Ministry of Natural Resources of China for providing the joint cruise, Dr. Zhibing Jiang and Dr. Dewang Li for analyzing total Chl a and inorganic nutrients samples and Dr. Cong Zeng for collecting the water samples. We are also thankful for Dr. Yisen Zhong for the CTD data generation and Shi Min for the flow cytometry technical support at Beckman Coulter International Trading (Shanghai) Co., Ltd.

## Supplementary Figures

**Fig. S1.**
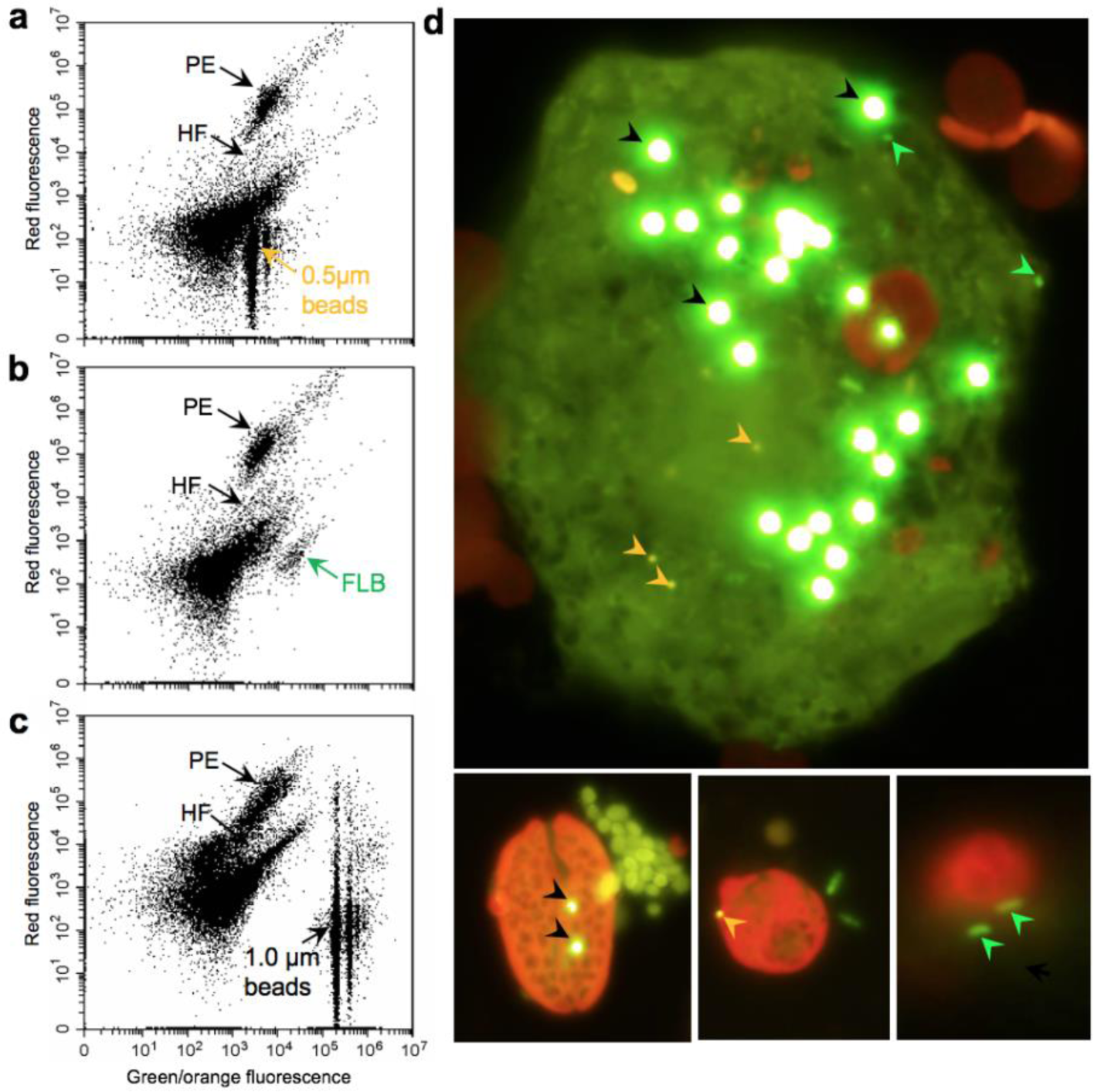
Comparison of identification results among different prey surrogates using both flow cytometry (**a-c**) and epifluorescence microscopy (**d**), including 0.5 µm yellow-orange fluorescent beads (**a**), fluorescent-labeled heat-killed bacteria, FLB (**b**), and 1.0 µm yellow-green fluorescent beads (**c**). Arrows indicate populations of pigmented eukaryotes (PE), heterotrophic flagellates (HF), and three different prey tested on flow cytometry in panel **a-c**. Panel **d** shows the intensity of excited prey fluorescent signals inside of a large bacterivore, marked with orange (0.5 µm beads), black (1.0 µm beads) and green arrow heads (FLB), respectively. Experiments were done prior to the cruise using lake samples.

**Fig. S2.**
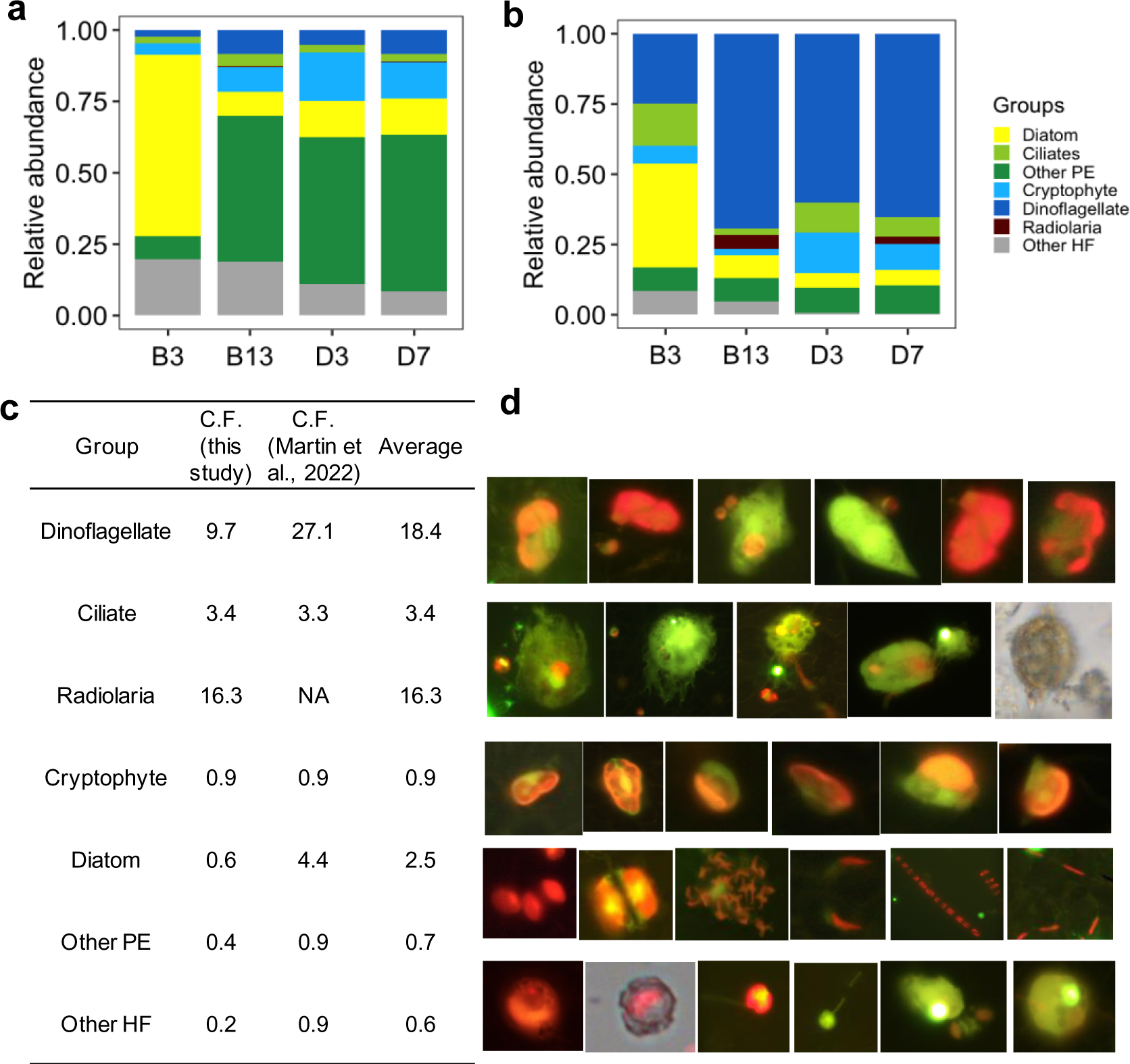
Comparison of relative abundances for seven distinguishable groups between cell abundances (microscopic observation) (**a**) and 18S rRNA gene abundances (**b**). The ratios between these two groups of results were used to generate our correction factors for getting 18S rRNA gene metabarcoding-derived cell abundances (**c**). Panel **d** demonstrate common and representable cellular morphology among those seven groups distinguished by microscopy, i.e. row one of dinoflagellates, row two of ciliates and radiolarian (the last image), row three of cryptophytes, row four of diatom, and row five of other PE (red fluorescent chloroplasts) and HF (no constitutive red fluorescent chloroplasts).

**Fig. S3.**
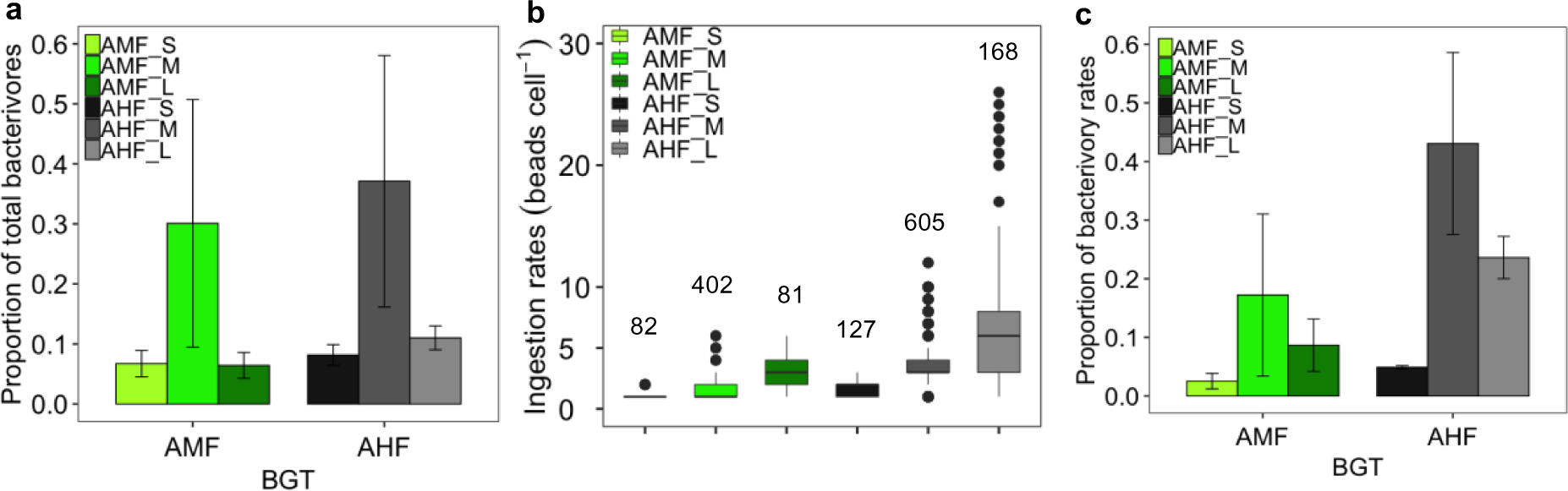
Averaged proportions in abundance (**a**), ingestion rates (**b**) and proportions of bacterivory contributions (**c**) of six binned bacterivore groups between active mixotrophs (AMF) and heterotrophs (AHF). All results were derived from epifluorescence microscopic observation and averaged among all four stations. Letters S, M, L in the legend denote small, medium and large sizes of <5 µm, 5-10 µm, and >10 µm, respectively, and numbers marked on top of each box in panel **b** denote numbers of cells identified for different groups.

**Fig. S4.**
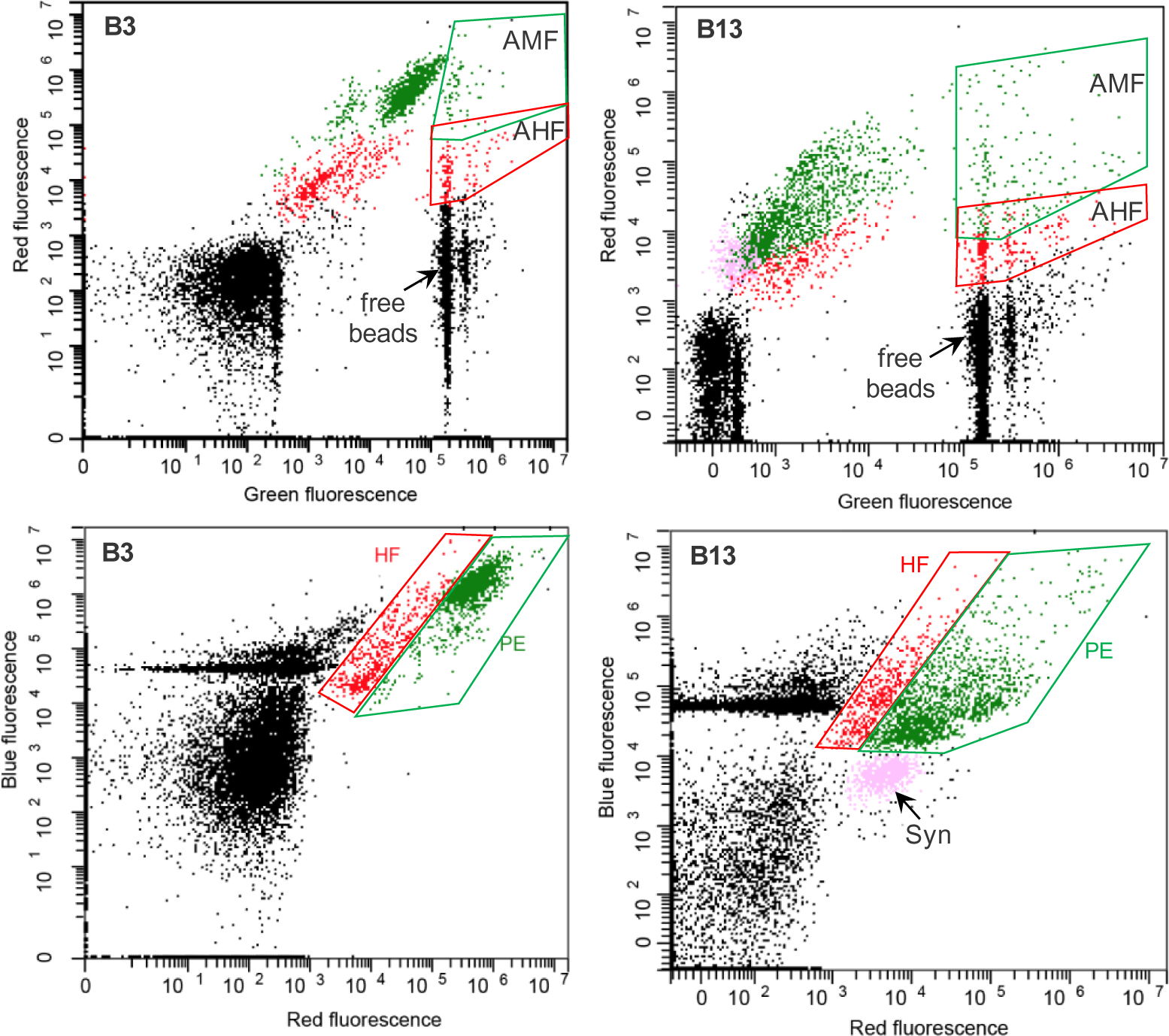
Representative flow cytometry plots from station B3 and B13 showing populations of pigmented eukaryotes (PE) in green dots and heterotrophic flagellates (HF) in red dots (with different fluorescent signals), distinguished initially based on red fluorescence (chlorophyll *a*) and blue fluorescence (DAPI stained nucleic acid) in lower panels (forward and side scatters were also check for cell sizes). Active bacterivores with high green fluorescence (prey tracer fluorescence) were separated from non-active eukaryotes, located on the right side of upper panel plots, including mixotrophs with higher red fluorescence (AMF) and heterotrophs (AHF) with lower red fluorescence. Another population of *Synechococcus* (Syn) was also identified from station B13 but not B3. Population clustering from stations D3 and D7 look similar as B13 on flow cytometry so those plots were not shown here.

**Fig. S5.**
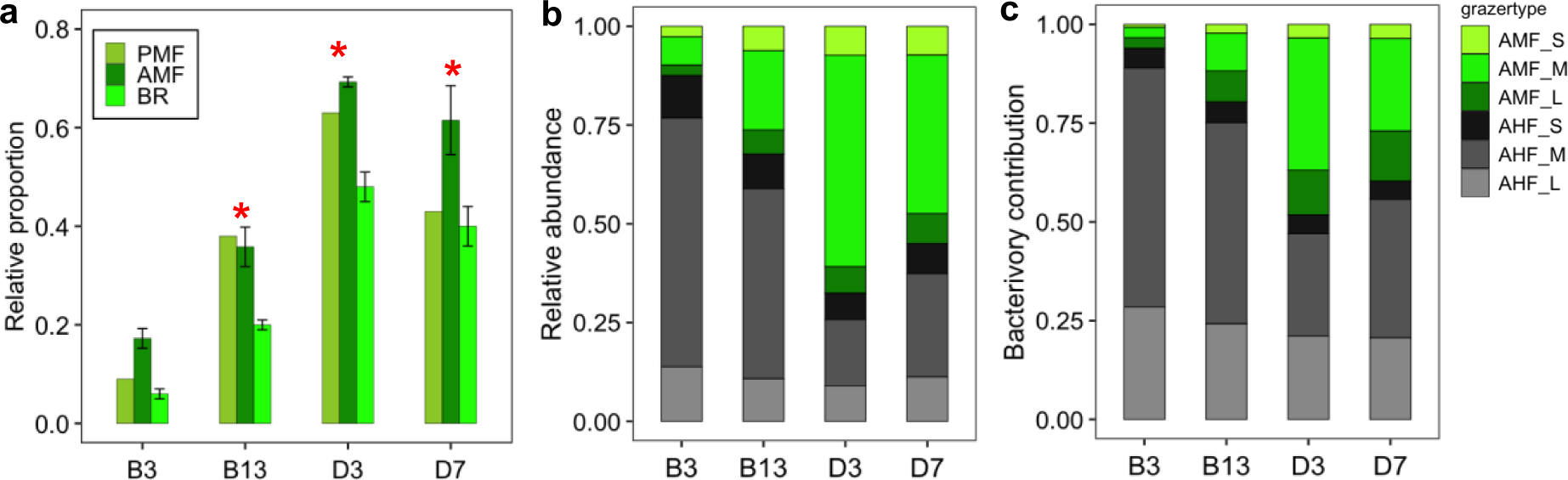
Panel **(a)** summarizes the prevalence of mixotrophs across stations using relative abundances of potential mixotrophs (PMF), active mixotrophs (AMF) and their bacterivory rates (BR). Asterisks indicate there was a significant difference between all three other stations against B3 among these three variables, based on pairwise student’s t-test (p<0.05). Panels **(b)** and (**c**) demonstrate realtive abundances and bacterivory contributions among different size groups of mixotrophs (AMF) and heterotrophs (AHF), retrieved from epifluorescence microscopic obervation.

**Fig. S6.**
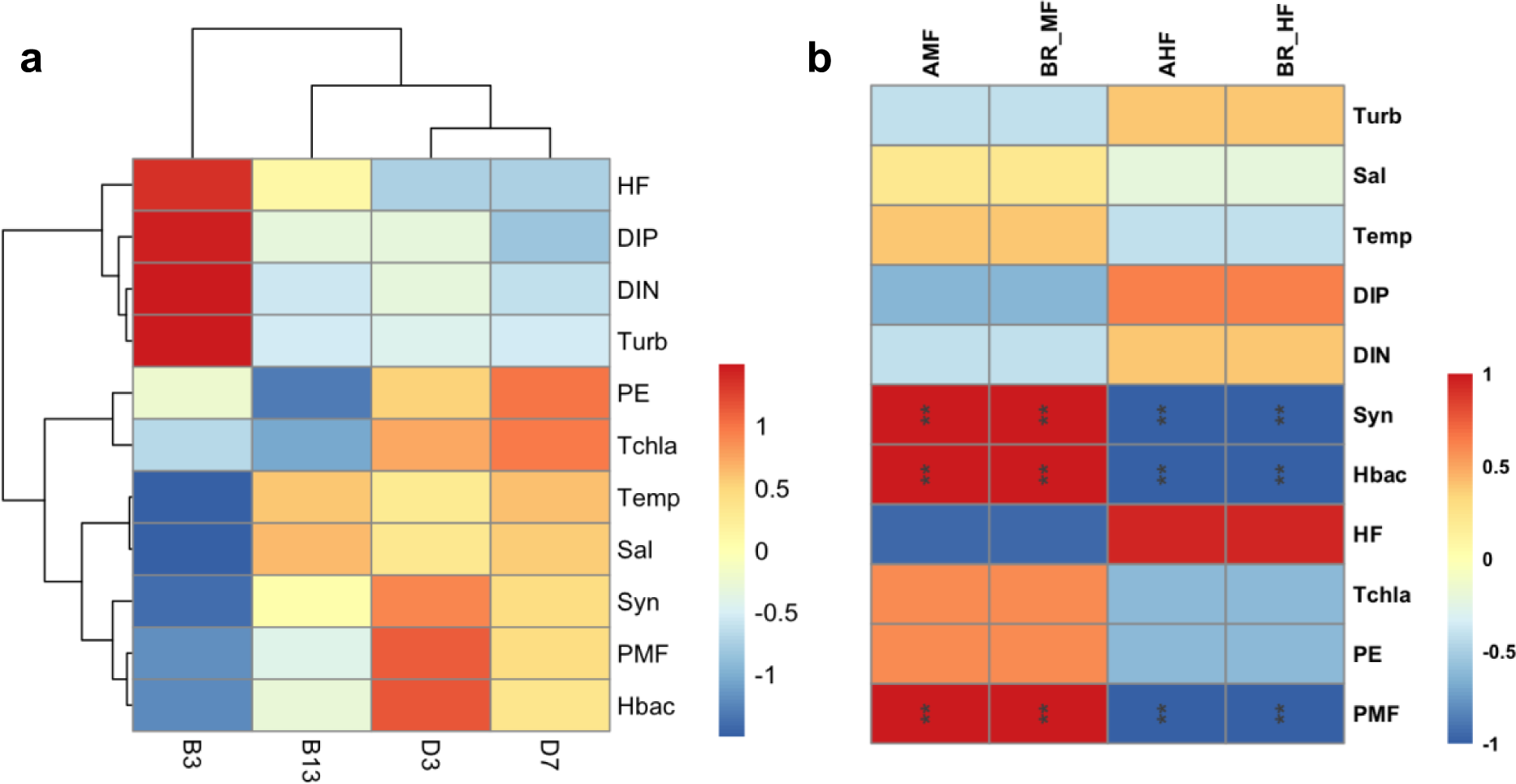
Panel (**a**) shows a heatmap of biological (HF of heterotrophic flagellates, PE of pigmented eukaryotes, Tchla of total chlorophyll *a*, Syn of *Synechococcus*, PMF of potential mixotrophs and Hbac of heterotrophic bacteria) and physiochemical variables (DIP, DIN, turbidity, temperature and salinity) across four stations. All variables were Z-score transformed (values shown in the color bar). Panel (**b**) presents the spearman correlation matrix between mixotrophic/heterotrophic fitness (AMF/AHF of active mixotrophs/heterotrophs and BR_MF/BR_HF of mixotrophic/heterotrophic bacterivory) and all independent variables shown in panel **a**. Asterisk marks reveals the significant correlations (P<0.05) and the color bar on the right denotes r values.

## Supporting information for active bacterivore identification based on flow cytometry method

Flow cytometry plots in the following figure (panel **a, b**) showing typical microbial populations of pigmented eukaryotes (PE), heterotrophic flagellates (HF) and cyanobacteria (cyan.), distinguished based on red fluorescence (chlorophyll *a*), blue fluorescence (DAPI stained nucleic acid) and green fluorescence (prey tracer fluorescence). PE and HF were initially gated based on blue and red fluorescence in panel **b** and each population was reflected on the plot in panel **a**, in green and red dots, respectively. Active bacterivores from PE and HF, on the right side of the plot, were identified based on high green fluorescence emitted from ingested prey (1.0 µm green-yellow fluorescent beads), and were further binned into population 1 (P1), population 2 (P2) and3 population 3 (P3). P2 was a small population (accounting for <10% of active mixotrophs or heterotrophs) located in between P1 and P3 with mixed mixotrophic and heterotrophic bacterivores. Three populations were confirmed by sorting and sequencing single cells from P1 (20 cells), P2 (7 cells) and P3 (7 cells), shown as a Bayes tree of 18S rRNA genes constructed with GTR substitution model and gamma variation rate in panel (**c**). Typical freshwater bacterivores (lake samples were used for this methodology validation purpose prior to the cruise) from Dictyochophyceae (mixotrophs) and Chrysophyceae (both mixotrophs and heterotrophs) were retrieved, with mixotrophs (genera *Pedinellales*, *Pseudopedinella*, environmental dictyochophyte clade, *Uroglena*, *Dinobryon*, and *Spiniferomonas*) from P1 (green color-shaded), heterotrophs (genus *Spumella*) from P3, and heterotrophs (genus *Paraphysomonas*) from P2 (red color-shaded). Fluorescent microscopic images in panel (**d**) show that putative mixotrophic dictyochophytes (left side) and heterotrophic chrysophytes (right side) were commonly found to ingest prey surrogates during the incubation. Gel electrophoresis of 18S rRNA gene bands in panel **e** show a high single cell genomic amplification efficiency (30/34) through multiple displacement amplification with the REPLI-g Single Cell Kit (Qiagen, USA), followed with a second PCR using specific 18S rRNA gene primers. Note sequence labels in panel **c** denote population_single cell number, and numbers in the bracket indicate cell numbers with the same taxonomy.

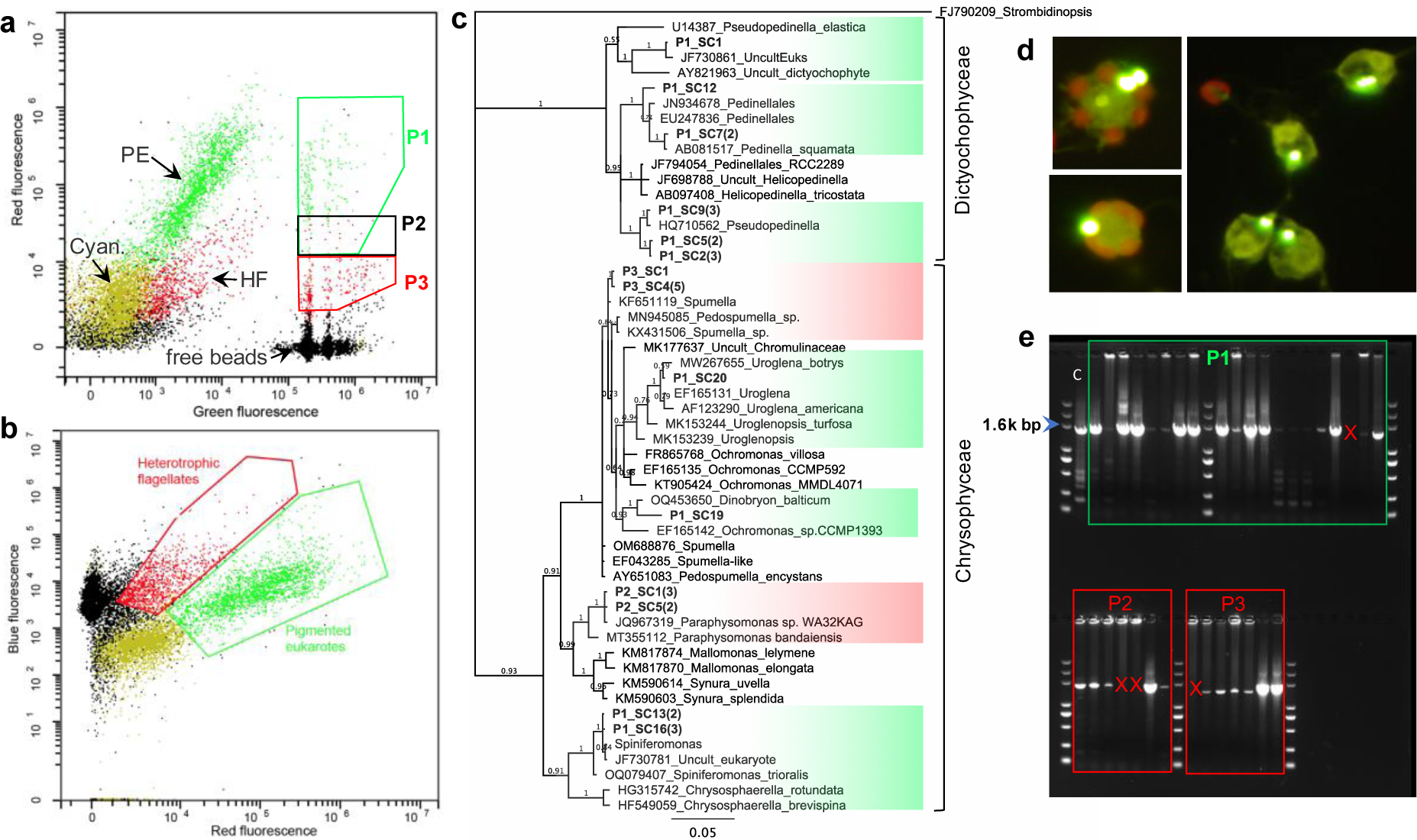

## Notes

### Competing Interest Statement

The authors have declared no competing interest.

